# Distinguishing Signal from Noise: Understanding Patterns of Non-Detections to Inform Accurate Quantitative Metabarcoding

**DOI:** 10.1101/2022.09.02.506420

**Authors:** Zachary Gold, Andrew Olaf Shelton, Helen R. Casendino, Joe Duprey, Ramón Gallego, Amy Van Cise, Mary Fisher, Alexander J. Jensen, Erin D’Agnese, Elizabeth Andruszkiewicz Allan, Ana Ramón-Laca, Maya Garber-Yonts, Michaela Labare, Kim M. Parsons, Ryan P. Kelly

**Affiliations:** Cooperative Institute for Climate, Ocean, & Ecosystem Studies, UW, Seattle, WA; Northwest Fisheries Science Center, NMFS/NOAA, Seattle, WA; School of Marine and Environmental Affairs, UW, Seattle, WA; School of Aquatic Fisheries Science, UW, Seattle, WA; Scripps Institution of Oceanography, UCSD, La Jolla

**Keywords:** quantitative metabarcoding, eDNA, amplicon sequencing, non-detection, zero inflation

## Abstract

Correcting for amplification biases in genetic metabarcoding data can yield quantitative estimates of template DNA concentrations. However, a major source of uncertainty in metabarcoding data is the presence of non-detections, where a technical PCR replicate fails to detect a species observed in other replicates. Such non-detections are an important special case of variability among technical replicates in metabarcoding data, particularly in environmental samples. While many sampling and amplification processes underlie observed variation in metabarcoding data, understanding the causes of non-detections is an important step in distinguishing signal from noise in metabarcoding studies. Here, we use both simulated and empirical data to 1) develop a qualitative understanding of how non-detections arise in metabarcoding data, 2) outline steps to recognize uninformative data in practice, and 3) identify the conditions under which amplicon sequence data can reliably detect underlying biological signals. We show in both simulations and empirical data that, for a given species, the rate of non-detections among technical replicates is a function of both the template DNA concentration and species-specific amplification efficiency. Consequently, we conclude metabarcoding datasets are strongly affected by (1) deterministic amplification biases during PCR and (2) stochastic sampling of amplicons during sequencing — both of which we can model — but also by (3) stochastic sampling of rare molecules prior to PCR, which remains a frontier for quantitative metabarcoding. Our results highlight the importance of estimating species-specific amplification efficiencies and critically evaluating patterns of non-detection in metabarcoding datasets to better distinguish environmental signal from the noise inherent in molecular detections of rare targets.

## Introduction

Metabarcoding, or DNA amplicon sequencing, is a powerful tool that can characterize biological communities without the need to physically observe individual organisms. The rise of metabarcoding via high-throughput sequencing is rapidly advancing human and wildlife health, ecology, and conservation science [1–8].

However, the power of metabarcoding applications lies in the ability to obtain reliable quantitative estimates of underlying communities [9–11]. In the case of metabarcoding and similar amplicon-based studies [12], it has become clear that 1) observations are non-linearly related to the underlying biology of interest [13,14], and 2) those observations are noisy, with many having relatively high variances as a function of expected values [11,15–17]. In order to make reliable quantitative estimates for any set of observations, we must be able to distinguish random variation from real signal. Thus, understanding the underlying signal-to-noise ratio is key to quantifying the power of detection for a given dataset [18].

Substantial efforts to correlate sequence reads and underlying community abundance have reported promising but largely equivocal results [10,15,19–23]. However, it is unsurprising that the application of simple linear correlations to non-linear and compositional datasets produce ambiguous results given the failure to model the underlying drivers of observed DNA sequence patterns and distributions. In response, recent mechanistic frameworks have begun to address the discrepancies between observed metabarcoding sequence counts and true underlying biological patterns by modeling the compounding processes that occur between DNA extraction and sequence observation [24–28]. These processes include DNA extraction, PCR, and multiple subsampling steps prior to sequencing [16,17,27,29,30]. Importantly, these mechanistic frameworks explicitly model the amplicon sequence-generating process by stating that observed sequence reads are a function of both the species-specific amplification efficiency and the underlying abundance of each species’ DNA within a sample [26]. Such models also reflect the inherent compositional nature of metabarcoding, acknowledging that metabarcoding data can only provide proportional (not absolute) abundances of a given species’ DNA in a given sample [11]. This approach can reconstruct starting DNA proportions, prior to PCR (e.g. [16,25–27]) and, where metabarcoding data are combined with additional information on underlying DNA concentrations, can yield absolute abundance estimates of the sampled DNA concentrations (e.g. [31]).

Despite these advances in modeling the amplicon sequence-generating process, it is clear that the sequential molecular steps required to generate metabarcoding data result in highly variable sequence-read counts among technical replicates derived from the same DNA extract [13,32–36]: replicate samples yield somewhat different results. Thus, in practice, it can be difficult to distinguish signal from noise in metabarcoding datasets. In particular, zeros are frequently over-represented in metabarcoding data, contributing substantially to among-replicate variability [16,24,34]. For example, in three technical replicates, a unique amplicon sequence variant (ASV) may be represented by 3,897; 165; and 0 reads across replicates (132,731, 196,260, 55,400 read depth for each replicate respectively; [31]). This observed variability among technical replicates far exceeds the expected variability arising from binomial- or multinomial sampling, and so demands a different explanation [16].

Here we focus on the patterns and causes of non-detections (in which a species is unobserved in one technical replicate despite being observed in other replicates) in metabarcoding datasets. After first synthesizing previous research on patterns of sequence counts, we simulate the process of metabarcoding to develop a qualitative understanding of the scenarios under which non-detections arise. We then use these results to generate predictions for the frequency of non-detections. Next, we use empirical observations to test these predictions using metabarcoding data derived from a set of ethanol-preserved fish larvae, in which both the underlying organismal abundances and the resulting metabarcoding dataset are well-characterized. Our empirical findings closely match the predictions and suggest a mechanism for non-detections and stochastic variability in general. Given this understanding of the sources of variability, we can more confidently distinguish signal from noise in metabarcoding datasets.

## Methods

### Conceptual Model and Simulating Metabarcoding Data

Our generating model for metabarcoding derives from Shelton et al. [27], building on the work of others [11,17,25,26,29]. Briefly, we envision a metabarcoding dataset as compositional, arising from a chain of sampling and amplification processes acting on individual DNA molecules.

We start with a sample of extracted DNA containing sequences from multiple species. From this starting point, there are many different metabarcoding laboratory protocols that lead to observed sequences from a sequencing instrument [37,38]. Here, we approximate this using three main stochastic processes following the commonly used two-step PCR library generation process (e.g., a target PCR followed by an indexing PCR). First, we assume a sample of DNA is extracted and included in the multi-taxon PCR reaction. Second, PCR amplification using a specific primer and protocol occurs, replicating the DNA molecules for each taxon. This second step includes the various target PCR, cleaning, indexing PCR, and pooling steps that occur during or following the main PCR reaction. Finally, the resulting mix of DNA copies is sampled to generate a compositional sample of amplicons that are observed through the sequencing instrument.

Mathematically, we can write a simulation for this framework as a series of linked stochastic processes. Specifically, we start with a sample of extracted DNA containing *λ_i_* copies *μL*^−1^ DNA from the *i*th taxon, *i = 1,2,…,I*. Let *W_ij_* be the discrete number of DNA molecules for species *i* sampled (i.e., in the tube in which a given PCR reaction takes place) of technical replicate *j* at the beginning of PCR and

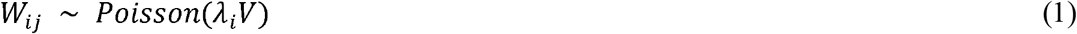

Here, *V* is the volume (*μL*) of template DNA sampled from the DNA extract. This equation assumes each taxon is sampled independently. Note that different technical replicates (e.g., *j* = 1 and *j* = 2) arise from the same environmental sample but may contain different numbers of molecules for a given species due to sampling variability.

Next, we model a three-step PCR process. Most importantly, we assume the amplicons produced during a PCR reaction are influenced by a species-specific amplification efficiency *a_i_*, which is characteristic of the interaction between the particular primer set, reaction chemistry, and template molecule of each species (*i*) being amplified [27]. For any species, *X_ij_* is the expected number of amplicons present in a technical replicate at the end of PCR. *X_ij_* is directly related to the efficiency of amplification and the starting number of DNA molecules, 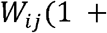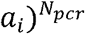, where *N_pcr_* is the number of PCR cycles and *a_i_* is bounded on (0,1); *a_i_* = 1 represents a perfect doubling of molecules with each PCR cycle. For the purpose of this paper, we assume a two-step PCR process with a sub-sampling and PCR cleaning process in between, and *X_ij_* can be modeled at each step as:

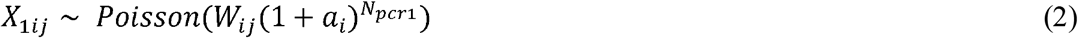

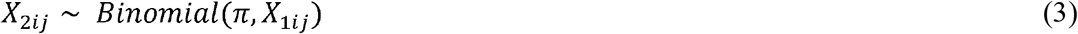

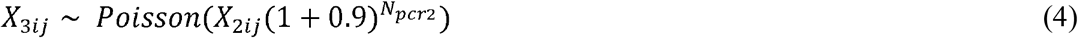

where *π* is the proportion of the first PCR product used in the second PCR amplification. Note that during the indexing reaction (equation 4) all taxa share a single amplification efficiency (*a_i_* =*0.9*) as we assume all indexing primers anneal to the indexing adapter sequences with equal efficiency. Here *X*_3_ is the number of amplicons present after both PCR amplifications but before sequencing. Finally, the sequencing instrument generates a total number of reads within technical replicate *j* (*N_readsj_*) and each replicate has a vector of observed read counts (***Y**_j_*, bolding indicates vectors) for *I* species.

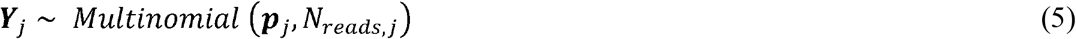

where *p* is the proportion of reads from species *i* in technical replicate *j*, and 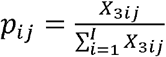. Thus, observed read counts (*Y_ij_*) are sampled stochastically based on their relative amplicon abundances, *p_ij_*.

The above model provides a general framework for understanding the causes of variability in observed read counts (*Y_ij_*), specifically the probability of non-detections as a factor of 1) the initial DNA concentration *λ_i_* and 2) the amount of species-specific variation in amplification efficiency (*a_i_*). The simulation allows us to identify two distinct causes of non-detection, *p*(*Y_ij_* = 0). First, non-detection may occur because there were no molecules of species *i* in the initial PCR (*W_ij_* = 0, in which case we are interested in *p*(*W_ij_* = 0) because *p*(*Y_ij_* = 0|*W_ij_* = 0) = 1). Second, zeros can arise due to PCR amplification and sequence-sampling processes; thus, we are interested in *p*(*Y_ij_* = 0|*W_ij_* > 0). While *p*(*W_ij_* = 0) is trivial to calculate from equation 1, determining *p*(*Y_ij_* = 0) and *p*(*Y_ij_* = 0|*W_ij_* > 0) are not. We turn to simulations to understand the contributions of variation in *λ_i_* and *a_i_* to the probability of non-detection.

We simulate four communities with different levels of richness (N = 4, 10, 30, and 50 taxa). For simplicity, we assume all taxa start with identical DNA concentrations regardless of the richness. DNA concentrations, *λ_i_* are varied from 0.5 to 10,000 copies *μL*^−1^ (for simplicity we set *V* = 1 for all simulations). We further allow a range of amplification efficiencies (*a_i_*) among taxa where *a* ~ *Beta*(0.7γ, 0.3γ) with γ ranging from 5 (high variation among species) to 1 million (no variation among species), but with a constant average amplification efficiency of 0.7 for all scenarios. We simulated 50,000 realizations for each combination of richness (4 levels), *λ* (18 levels), and *γ* (6 levels: 5, 10, 20, 100, 100, 1 million)), for a total of 432 scenarios. For all the simulations, we allowed sequencing depth to vary among replicates (*N_read,j_* was uniformly drawn from discrete values between 60,000 and 140,000), used a fixed sampling fraction (*π* = 0.20), and set *N_pcr2_* = 35 and *N_pcr2_* = 10). We calculated a range of summary statistics for each scenario, including the overall probability of non-detection, *p*(*Y_ij_* = 0); the probability of non-detection due to the absence of the target molecule, *p*(*W_ij_* = 0); and summaries of the reads both in absolute terms and in terms of relative abundance.

#### Empirical Testing

We test these hypotheses against a real metabarcoding data set making use of three data streams generated from a common set of biological samples: organismal abundance (as a proxy for input DNA molecules), metabarcoding data, and amplification-efficiency estimates for the relevant species.

### Study Design

As part of the California Cooperative Oceanic Fisheries Investigations (CalCOFI), Gold et al. [31] use morphological and molecular methods to analyze the response of ichthyoplankton in the California Current Large Marine Ecosystem to ocean warming. Ichthyoplankton samples were collected in oblique bongo net tows on CalCOFI research cruises over two decades (1996; 1998-2019). Once a sampling tow concluded, ichthyoplankton present on one side of the net were preserved in Tris-buffered 95% ethanol and stored in the Pelagic Invertebrate Collection at Scripps Institution of Oceanography [39]. The paired ichthyoplankton were preserved in sodium borate-buffered 2% formaldehyde for microscopy-derived species identification and abundance (number of larvae per species per jar). This dataset yields paired samples for both metabarcoding analysis and absolute abundance counts from the same sampling event.

### Abundance Estimation from Microscopy

Formalin-preserved larvae were identified and enumerated following the methods of Thompson et al. [40]. The majority of taxa were identified to species level. Here we assume that the relationship of absolute abundance (counts of individual species) is proportional to the amount of species-specific DNA in the extraction. See the discussion for the merits of this assumption.

### Metabarcoding Data Generation

DNA sequences were generated from 84 ethanol-preserved samples as described in Gold et al. [31]. Briefly, ethanol samples were filtered onto 0.2 μm PVDF filters and were extracted using a Qiagen DNeasy Blood and Tissue kit. We then amplified three technical PCR replicates using a touchdown PCR and the MiFish Universal Teleost specific primer [41]. Both a negative control (molecular grade water instead of DNA extract) and two positive controls (DNA extract from non-native, non-target species) were included alongside samples. Libraries were prepared using Illumina Nextera indices following the methods of Curd et al. [42] and sequenced on a NextSeq 2x 150 bp mid output. Sequencing data was then processed using the *Anacapa Toolkit* [42] to conduct quality control, ASV dereplication, and taxonomic assignment. Sequences were annotated with the California fish specific reference database and a bootstrap confidence cutoff score of 60 following the methods of Gold et al. [43]. Eight technical replicates with either low sequencing depth (n<30,000) or high dissimilarity (Bray Curtis dissimilarity > 0.7) were removed.

### Amplification Efficiency Estimation from Mock Communities

We used a subset of the mock communities generated for Shelton et al. [27] to estimate amplification efficiencies of relevant fish species. Mock communities included DNA from 57 voucher fish tissue samples, 17 of which were detected in the CalCOFI metabarcoding data set, from the Scripps Institution of Oceanography Marine Vertebrate Collection. To accurately quantify input DNA for each species within the mock community, we used a nested PCR strategy in which mock communities were generated by pooling resultant longer fragment PCR products of each species rather than by pooling the total genomic DNA of each species (which includes variable amounts of nDNA as well as bacterial and other DNA sources). To implement our nested PCR strategy, we first amplified a 612 bp fragment of the *12S* rRNA gene that contains the MiFish Universal Teleost *12S* primer set [44], and quantified the resulting PCR products using the QuBit Broad Range dsDNA assay (Thermofisher Scientific, Inc.); this yielded measurements of species-specific, amplifiable DNA. Using this known-concentration DNA we generated 9 distinct mock communities by pooling long fragment PCR products comprising three distinct sets of species and three abundance distributions (See Table S1). Pooled mock communities were using the QuBit Broad Range dsDNA assay (estimated concentrations ranged from 8-12 ng μL^−1^) and then diluted serially by a 1:10 dilution down to 10^−8^ original concentration. We then converted ng μL^−1^ to copies μL^−1^ using the following equation:

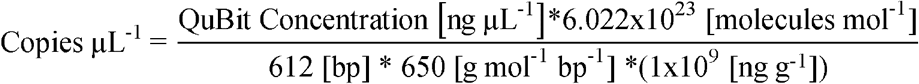

Finally, input concentrations of 380-600 DNA copies μL^−1^ for each total community were loaded in the MiFish Universal Teleost *12S* PCR step (Table S2). Given this design, each mock community had a different number of DNA molecules per species. We then amplified each of the mock communities in triplicate with the MiFish Universal Teleost *12S* following the methods of Curd et al. [42], targeting a 185 bp fragment within the larger 612 bp PCR fragment used to generate the mock communities. Each triplicate PCR technical replicate was then treated as a unique library and sequenced separately. Metabarcoding libraries were then prepared and sequenced on a MiSeq platform using a v3 600 cartridge following the methods of Gold et al. [43]. We note that one set of mock communities were re-sequenced on a separate run to generate usable data. Resulting sequences were processed using the *Anacapa Toolkit* using the global *CRUX* generated reference database given the broad geographic distribution of species from Gold et al. [43]. We also used a taxonomic cutoff score of 60 as above. Taxonomic assignment of ASVs was confirmed with BLAST using default settings. For the two observed discrepancies, we chose to use BLAST assignments with greater than 99% identity and 100% query length match as they matched our known vouchered specimen identifications.

We fit the model from Shelton et al. [27] to a third of the data (3 technical replicates of each evenly pooled mock community). Generated parameter estimates were then used to predict the starting proportions of DNA in the remaining two-thirds of the data, for an out-of-sample estimate of accuracy. We used the resulting model output to calculate the mean amplification efficiency per species. The model, implementation, and code are detailed in Shelton et al. [27], but there are two particularly relevant points from the model for connecting the simulation and empirical results that we highlight here. While we simulate absolute amplification efficiencies (*a_i_*), because metabarcoding data is compositional, the absolute amplification efficiency cannot be estimated from metabarcoding data. Instead, we estimate amplification efficiencies for each species relative to a reference efficiency (see also [25,26]). In our case we estimate *α_i_* as the amplification efficiency of species *i*, *a_i_*, relative to the efficiency of a reference species, *a_R_*, therefore 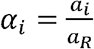. Thus, for simulations we discuss *a* but for estimation, we discuss *α*. Note while values of *α* can be directly calculated from *a*, values of *a* are not uniquely identifiable from *α*.

All data and code for conducting analyses will be made publicly available upon acceptance via NCBI SRA, Dryad, and GitHub (https://github.com/zjgold/Metabarcodings_Signal_from_Noise).

### Hypothesis Testing

From the above outlined simulations and empirical data, we generate a series of hypotheses. First, we expect fewer non-detections for more abundant DNA molecules, given the same species (and therefore the same amplification efficiency). Second, we hypothesize that species with higher amplification efficiencies will have fewer non-detections and higher observed sequence read counts than species with lower amplification efficiencies, given the same abundance of template DNA molecules. Third, we expect species with low amplification efficiencies will have a high rate of non-detection regardless of the abundance of template DNA molecules. We test each of these hypotheses using both simulation and empirical results.

## Results

### Simulation Results

We found a strong correlation between the probability of non-detections and both the absolute abundance of template DNA molecules and amplification efficiencies (Figure 1). The probability of non-detections (*p(Y=0)*) dramatically declines when concentrations of template DNA are greater than ~10 copies μL^−1^ per species, given an average amplification efficiency of 0.7 (Figures 1C & 1D). Likewise, our results demonstrate that species with low amplification efficiencies exhibit high probabilities of non-detections regardless of starting DNA concentrations (Figure 1A, B). Importantly, we demonstrate that even species with an amplification efficiency slightly below average (e.g., *a* = 0.7) exhibit high rates of non-detections at DNA concentrations far higher than from typical eDNA field samples (e.g. *λ* > 100 copies/μL; [45]). Together these simulations indicate that the probability of non-detection is dominated by the subsampling process at low template DNA concentrations while the probability of non-detection is driven primarily by the PCR process (i.e., differences in amplification efficiencies) at higher template DNA concentrations.

**Figure 1.**
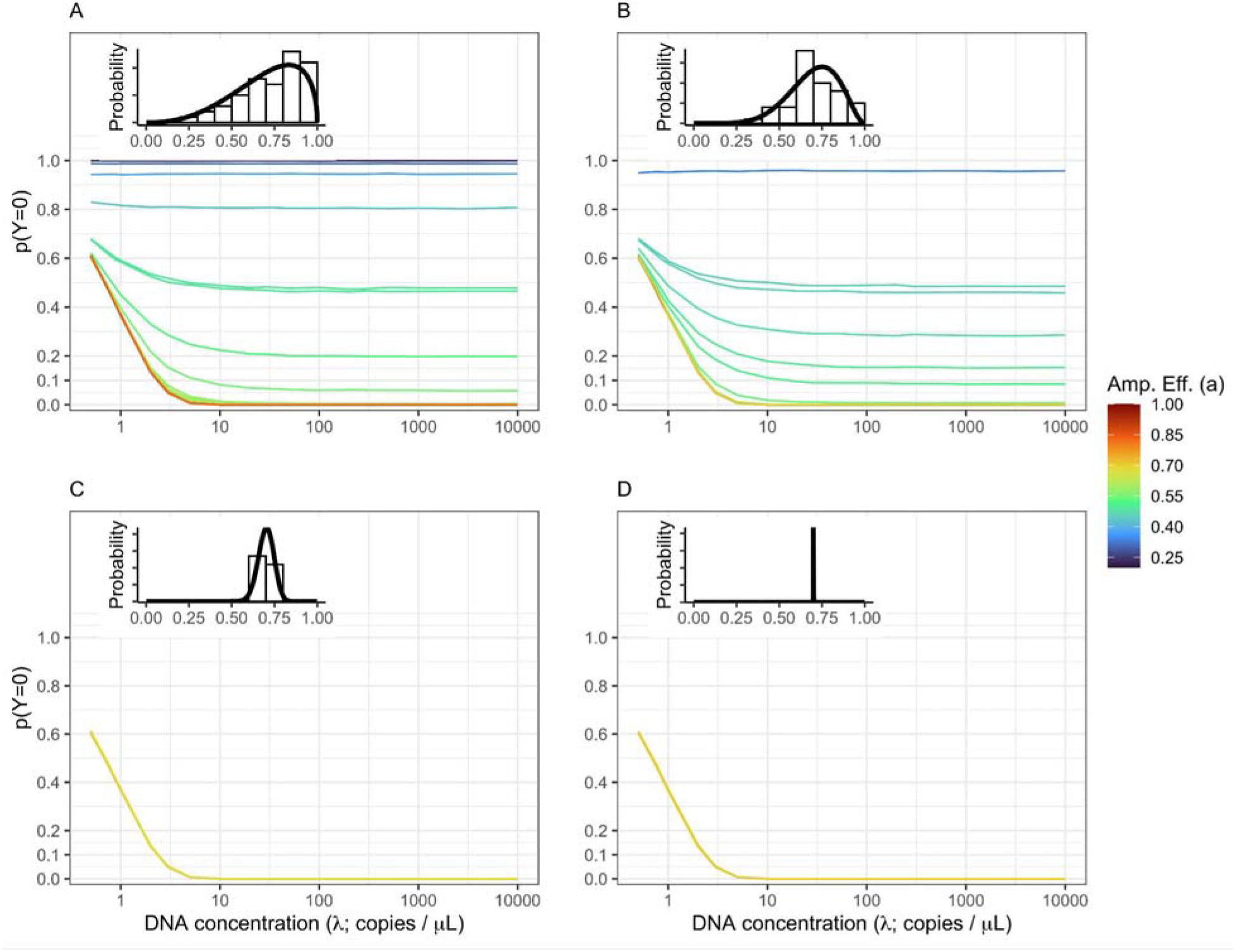
Non-detections Driven By Both DNA Concentration and Amplification Efficiency. The probability of non-detection (*p(Y=0)*) is shown for a community of 50 equally abundant taxa with the amplification efficiency distribution shown inset in the upper left of each panel. The amount of among-taxa variation in amplification efficiency varies from high variation *(A;* γ=5) to moderate variation (*B*: γ=10) to low variation (*C*: γ=100) to effectively no variation (*D*: γ=1,000,000). Both subsampling and amplification efficiencies influence the rate of non-detection. The probability of observing no DNA in a given technical replicate is highest at low DNA concentrations (<10 copies /μL). However, non-detections are possible for species with below average amplification efficiencies (in this case approximately *a_i_* = 0.7) and very likely (*p(Y=0)* > 0.5) for amplification well below average (*a_i_* < *0.4)*.

### Empirical Results

#### Microscopy Results

Independent estimates of abundance were generated from sorting 9,610 larvae from 84 jars (min = 2, max =960). See Gold et al. [31] for a detailed description of the results.

#### Metabarcoding Results

The metabarcoding data set generated from ethanol-derived eDNA consisted of a total of 54.5 million amplicon sequence reads that passed through the *Anacapa Toolkit* quality control, ASV dereplication, and decontamination processes. Sequencing depth ranged from 36,050 reads to 1.2 million reads per technical replicate. For our integrated Bayesian model of the probability of non-detection in a technical replicate, we focused on the 17 species that had 1) sufficient representation across the metabarcoding data set (observed in > 10 technical PCR replicates) to achieve model convergence and 2) were represented in our mock communities. See Gold et al. [31] for the full description of model implementation and results.

#### Mock Community Results

The mock community data set consisted of 4.0 million amplicon sequence reads that passed through the *Anacapa Toolkit* quality control, ASV dereplication, and decontamination processes across a total of 36 unique samples comprising three distinct community assemblages each with three PCR technical replicates. Sequencing depth ranged from 9,872 reads to 206,900 reads per technical replicate. Of the 57 voucher species represented, we classified 56 unique species, and used the Shelton et al. [27] model to estimate amplification efficiencies for each species. One species, *Urobatis halleri*, was not detected in any technical replicate. *Citharichthys sordidus* was present in all mock communities and was selected as the reference species for estimating relative amplification efficiencies. Across all species, *α_i_* ranged from −0.30 to 0.03 with a mean of −0.06 (Table S3). For presentation purposes, we label species with *α_i_* values below −0.07 as a low amplification efficiency group (n=15) and the remaining species as a high amplification (n=41).

### Hypothesis Testing

As with the simulation results, we found that the probability of non-detections is strongly correlated with both the abundance of DNA molecules for a given species within a sample and the species-specific amplification efficiency (Figures 2b, 3b). Non-detections occur more frequently at low DNA concentrations regardless of amplification efficiency (Figures 2b, 3b). Species exhibiting lower amplification efficiencies (*α_i_* < −0.07) had higher rates of non-detections even at high input DNA concentrations (10^4^ copies μL^−1^) and larval counts (9 larvae per jar; Figures 2b, 3b).

**Figure 2:**
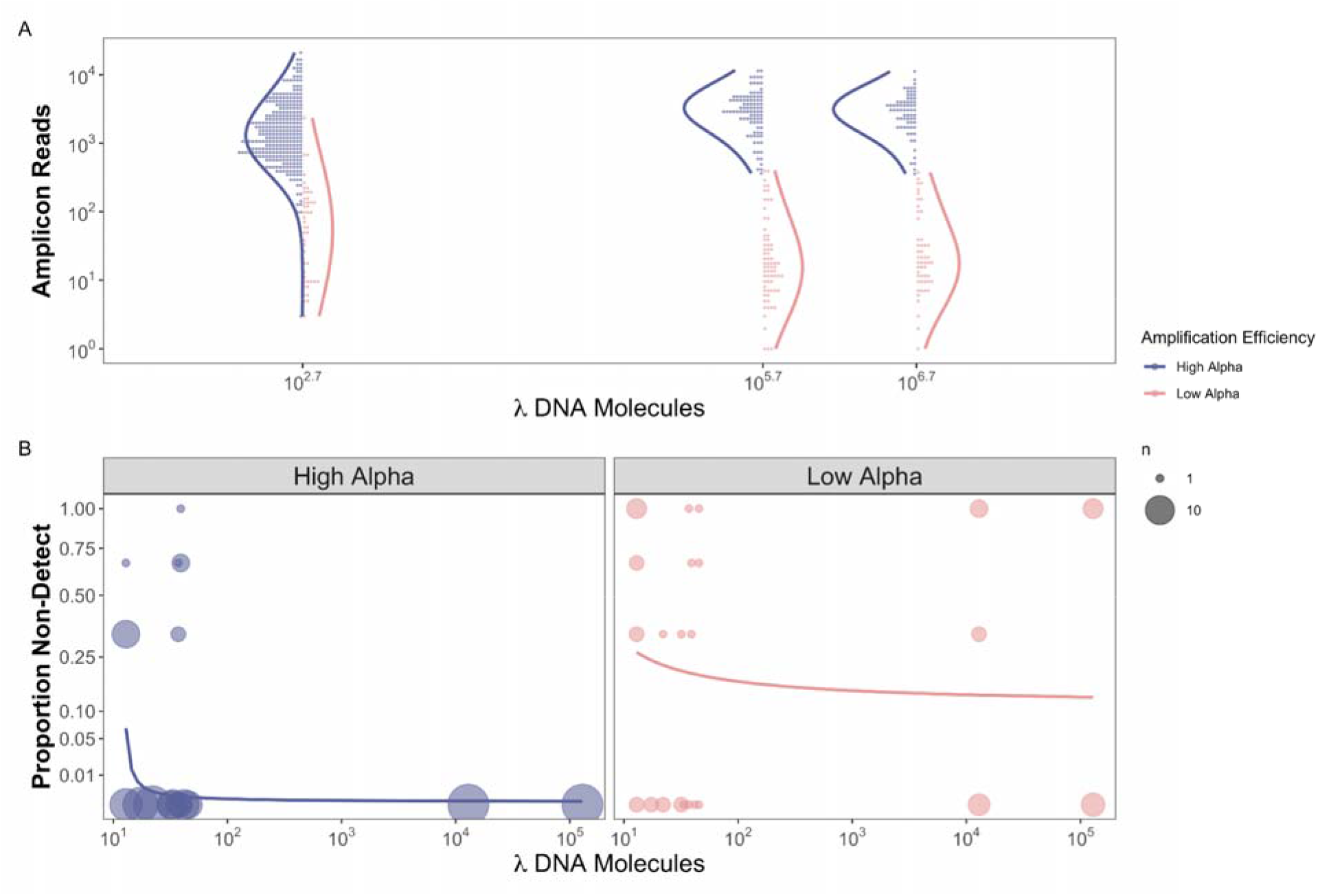
Observed Reads and Non-detections are a function of Amplification Efficiency and Input DNA Concentration in the Mock Community example. For species observed within a replicate, we find that species with higher amplification efficiencies (>0.7) have a greater number of observed reads for an equivalent template DNA concentration (a). We also find no difference in the total number of observed reads and increased DNA concentration as expected for a compositional data set. Furthermore, we find a greater proportion of non-detections when both DNA concentration and amplification efficiencies are lower (b). These results align well with our simulated data.

Furthermore, from the mock community example, species with higher amplification efficiencies (*α_i_* >−0.07) have higher observed sequence read counts for an equivalent template DNA concentration (Figure 2a). The 41 species with high amplification efficiencies have more reads sequenced per DNA molecule added (mean ± sd = 4.1 ± 6.31, range = 0.00-55.4) than the 15 species with low amplification efficiencies (mean ± sd = 0.1 ± 0.47, range = 0.00-5.5). Likewise, species with higher larval counts in ethanol-preserved samples from plankton tows also have higher observed sequence read counts (Figure 3a). From the CalCOFI example, the 15 species with high amplification efficiencies have more reads sequenced per larvae counted (mean ± sd = 6,689 ± 28,305, range = 0-79,454) than the two species in the low amplification efficiency group (mean ± sd = 524 ± 1,080, range = 0-7,101).

**Figure 3.**
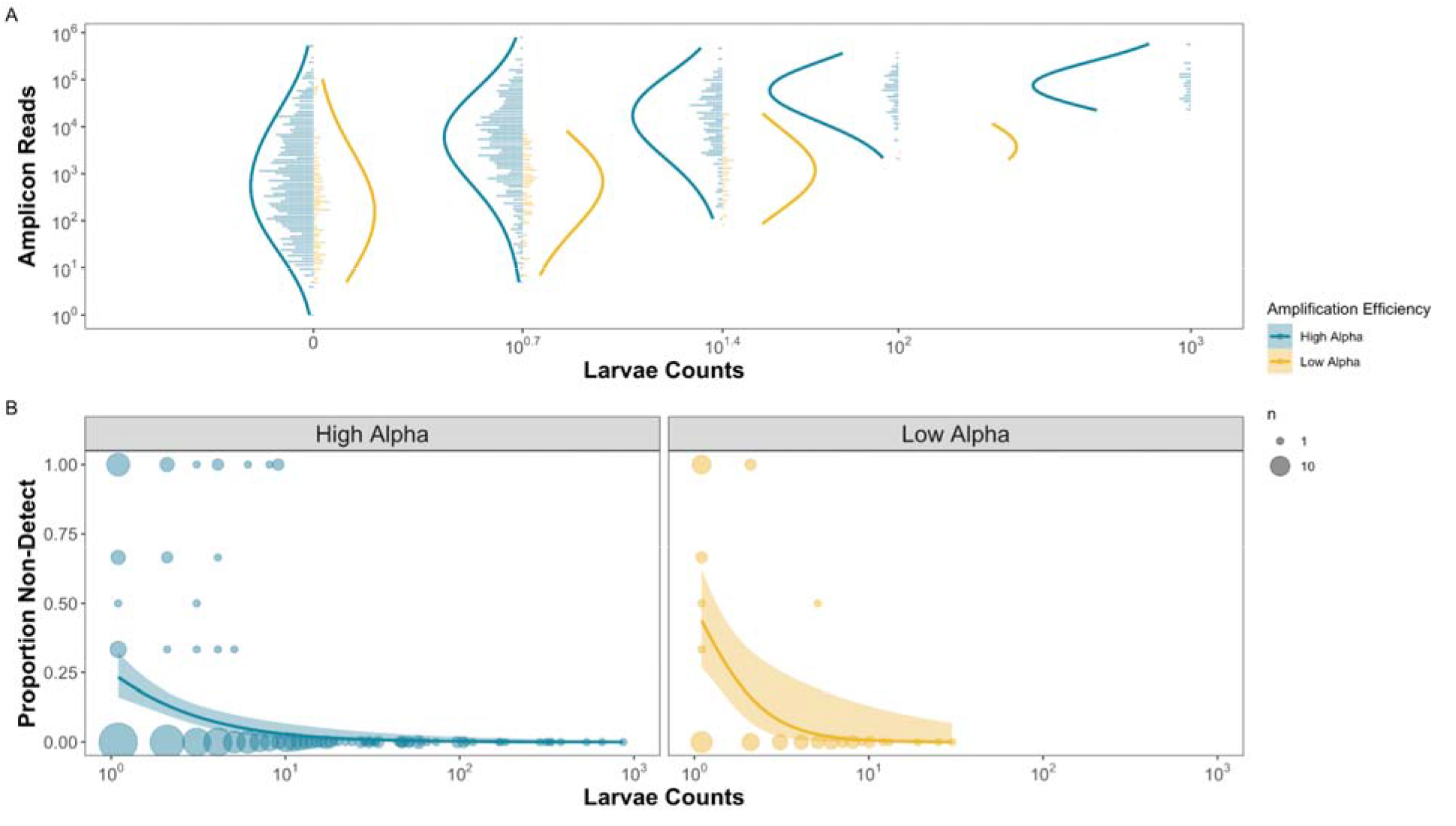
Observed Reads and Non-detections are a function of Amplification Efficiency and Larval Abundances in the CalCOFI example. For species observed within a replicate, we find that species with higher amplification efficiencies (>−0.07) have consistently greater numbers of observed reads for an equivalent template DNA concentration (a). We assume that the number of larvae in the jar is proportional to the number of DNA molecules present. We also find a greater proportion of non-detections when larvae are rare in the jars and have lower amplification efficiencies (b).

## Discussion

Using both simulated and empirical data, we demonstrate that observed sequence read counts from metabarcoding data are a function of species-specific input DNA concentrations, subsampling, and species-specific amplification efficiencies. Variability among replicates in detections of specific taxa – reflecting either rare targets or poor amplification efficiencies – are a substantial source of noise in these data. Consequently, it can be difficult to distinguish signal from noise in metabarcoding datasets. Our results illustrate several potential causes of non-detections and suggest that metabarcoding data can provide reliable quantitative estimates for species with abundant input DNA (> ~50 copies μL^−1^) and high species-specific amplification efficiencies. By characterizing underlying sources of sequence read count variability in metabarcoding, we identify key sources of noise that impact our ability to derive quantitative estimates of source DNA.

### Subsampling Rare Targets Results in Non-detections

Consistent with expectation, our framework strongly suggests that all else being equal in a metabarcoding assay (e.g., assuming even amplification efficiencies across species), rarer template DNA molecules have a higher probability of non-detection across technical replicates. These findings align well with observations of qPCR assays in which the probability of non-detection increases as you approach the limit of detection, in terms of absolute copies of DNA per reaction volume [46,47]. High rates of non-detections in qPCR assays are commonly observed for input DNA concentrations between 1 and 10 copies [46,48,49] and are likely driven by subsampling errors in which too few or no physical DNA molecules are transferred into a given PCR reaction [36,50,51]. These observations from qPCR studies reflect the findings from both simulated and empirical metabarcoding results reported here.

Importantly, subsampling rare target DNA molecules yields stark differences in observed per-species read counts among technical replicates, non-detections being the most obvious case of this phenomenon [15,16,34,46]. Together, these findings strongly support the hypothesis that the concentration of target DNA within a sample influences the observed patterns of amplicon read counts, particularly increasing the probability of non-detections for species with low template DNA concentrations. Such observations of high rates of non-detections also justify the use of over dispersed multinomial sampling approaches within metabarcoding models [27].

### Amplification Efficiencies Drive Sequence Counts and Non-detections

While the relationship between template concentration and non-detections is well documented in the literature [10,12,16,24,34,46], the causes of non-detection among species with abundant DNA are not widely appreciated. Both simulation and empirical results demonstrate that species with higher amplification efficiencies have higher observed amplicon read counts, confirming the predictions of previous compositional modeling efforts [25–27]. Furthermore, we find a clear association between the probability of non-detections and amplification efficiencies, with species with higher amplification efficiencies exhibiting fewer non-detections. Here we observed an order of magnitude difference in average sequence reads per larvae collected in a given jar with a maximum observed difference in amplification efficiencies of 0.33 (n=17). Previous research using mock communities has similarly demonstrated that equal concentrations of DNA in a single extraction frequently results in amplicon read counts that differ by orders of magnitude [16,32,36,37,52]. Such dramatic differences in resulting read proportions are understandable given the exponential nature of PCR - even a subtle difference in amplification efficiency across 30+ PCR cycles can result in stark differences in sequence counts [27].

The observed variation in amplification efficiency among species in metabarcoding approaches arises from complex PCR processes, including primer specificity, DNA polymerase selectivity, annealing temperature, GC content, and higher-order dimensional structure of DNA, inhibition, and co-factors such as MgCl_2_, among others [53–59]. This complexity makes designing metabarcoding assays that are highly specific for only target taxa challenging [60,61], resulting in the amplification of off-target taxa as well as a range of amplification efficiencies across target taxa [26,27,43,62,63]. As demonstrated by our simulations and empirical results, such a range of amplification efficiencies can result in substantial noise in metabarcoding data sets.

### Complex Relationship between Amplification Efficiencies and Abundance

The above results highlight the cumulative importance of the variance in amplification efficiency among species, as well as the abundance of template DNA for understanding the patterns of metabarcoding non-detections. The interaction between these factors is key for disentangling the signal from the noise of metabarcoding data. Here, we demonstrate that there are two ways to obtain non-detections for a given species after sequencing: low initial DNA concentration or low amplification efficiency. Both of these results are clear from our empirical CalCOFI fish larvae dataset which captured the effects of species-specific amplification efficiency and DNA concentrations on both sequence read counts and frequency of non-detections (Figure 2). Importantly, our results demonstrate that noise in metabarcoding datasets, like signal, is non-random and can be accounted for [16].

Alone, metabarcoding data is insufficient to tease apart these complex interactions. However, distinguishing signal from noise in metabarcoding datasets is tractable using independent estimates of amplification efficiencies and underlying DNA concentrations. Amplification efficiencies can be estimated through either generating mock communities [26,27], by amplifying a subset of samples multiple times at various numbers of PCR cycles [25], or by including internal positive controls within each PCR [28]. Likewise, underlying DNA concentrations can be estimated using qPCR or dPCR assays of key taxa or the metabarcoding locus itself; or estimated using non-genetic independent abundance estimates such as the microscopy counts presented above. As demonstrated here, and in Shelton et al. [27], McLaren et al. [26], and Silverman et al. [25], the inclusion of independent estimates of amplification efficiencies and DNA concentrations allow for the delineation of signal from noise from metabarcoding data sets. Further modeling efforts incorporating stochastic sampling of rare molecules prior to PCR will allow for accurate quantification and identification of true absences in metabarcoding data sets, greatly enhancing biological and ecological interpretation.

Furthermore, our analysis also underscores the importance of technical PCR replicates to quantify sequence variance in metabarcoding studies [64–66]. Without technical replicates, we would not have been able to quantify the frequency of non-detections in our metabarcoding datasets [17]. We demonstrate that non-detections may indicate low-relative-abundance starting DNA concentrations regardless of observed read depth, and conversely, may indicate low amplification efficiency regardless of starting concentration [27]. Thus, our results strongly support the inclusion of technical replicates for metabarcoding studies, particularly for deriving quantitative estimates.

Current best practices for qPCR and dPCR assays include numerous technical replicates to help distinguish signal from noise [46,48]. However, we recognize that technical replication dramatically increases the cost and effort of metabarcoding projects and may exhaust limited DNA extracts and resources. Alternatively, technical replicates could be performed on a subset of samples and the observed variance could be used to contextualize sequence read patterns in the whole dataset. However, such approaches come with a suite of assumptions, particularly whether the pattern of species’ sequence counts behaves similarly across all samples and environments/treatments. Future efforts to validate such approaches are clearly warranted.

In addition, given the importance of subsampling in driving non-detections, our results strongly suggest that field and laboratory processes that increase the absolute abundance of DNA molecules will reduce the noise in observed amplicon sequence reads [67]. For example, using a greater volume of DNA template for PCR reactions (3 μL vs. 1 μL) will reduce subsampling driven non-detections across samples. Likewise, increasing the total amount of water filtered for eDNA samples (3 L vs. 1 L) acts to concentrate DNA from the environment, similarly reducing subsampling driven non-detections [68]. These are two of many examples of laboratory protocols that may serve to increase the available number of DNA molecules and reduce the impacts of subsampling rare molecules, consequently improving quantitative estimates from amplicon sequence data.

The above mechanistic frameworks focus on processes from DNA extraction through sequencing, but do not approach the myriad of factors that influence the amount of DNA collected from the environment, gut, or other starting communities for metabarcoding. Substantial efforts have focused on understanding the effects of gene copy number, patchiness, shedding and degradation rates, and the fate and transport of cellular DNA, among others, on the amount/types of DNA collected from the environment [26,51,69]. Linking such research to the growing body of work that quantifies sources of potential bias in the lab, including the present study, is an important next step in understanding the relationship between biological signals and observed sequence read counts.

We recognize that incorporating the additional laboratory analyses and technical replicates to better characterize metabarcoding results may not be feasible for all metabarcoding applications. Many metabarcoding efforts are exploratory in nature, primarily focused on the characterization of biodiversity in under sampled habitats including the deep sea, polar regions, remote alpine regions, etc. For such exploratory biodiversity surveys, the additional efforts needed to achieve quantitative metabarcoding outlined above may not be practicable given surveying and budget constraints. However, it is important to recognize that our framework extends not only to quantitative metabarcoding but detection rates of taxa from metabarcoding surveys. The expected detection rate (observed reads > 0) of a given taxon in metabarcoding data is a function of other species in the community, the amplification rate of the target species, the amplification rates of other species, the proportional abundance of the target species, and the absolute abundance of the target species as demonstrated in our empirical datasets above. Thus, estimating the probability of detection from metabarcoding data alone is difficult in the abstract, but is quite tractable given a set of estimated parameters for a particular sampled community. Conversely, interpreting metabarcoding results from exploratory applications within systems with limited ecological context is challenging as species detection rates are a function of multiple unsampled parameters.

Undoubtedly, addressing this shortcoming of compositional metabarcoding data requires increased field and laboratory efforts. Such challenges are acute in under studied systems where the creation of mock communities is particularly difficult with limited access to vouchered DNA samples, let alone known species lists. However, exploratory metabarcoding studies do not preclude the revisiting of quantitative metabarcoding approaches in the future, especially since DNA extracts can be archived. For example, metabarcoding data can be generated first to provide an initial perspective into community assemblages that then allows for the identification and development of single species qPCR/dPCR assays and mock communities or variable PCR targets. In summary, we argue that all future best practices of metabarcoding results incorporate additional independent estimates of amplification efficiency, independent estimates of DNA concentrations, and technical replicates to better contextualize metabarcoding efforts. Given the rapid decline in sequencing costs and steady improvement in the development and implementation of molecular assays, such additional work is tractable, opening the door to adoption for routine application across metabarcoding studies to generate characterization of underlying biological communities.

## Conclusion

Ultimately, we demonstrate that variation in amplification efficiencies and underlying template DNA concentration are responsible for a substantial portion of observed noise in metabarcoding datasets. This study demonstrates the value of incorporating additional independent estimates of amplification efficiencies and DNA concentration along with amplicon sequence data, providing for the application of routine statistical approaches and straightforward interpretation of observed read patterns. Together with Shelton et al. [27], we provide a framework for establishing reliable estimates of abundance from amplicon sequence data that will be critical for extending the application of this method to health and ecological questions.

## Data Availability Statement

All data and code for analyses will be made publicly available at NCBI SRA, Dryad, and Github (https://github.com/zjgold/Metabarcodings_Signal_from_Noise) upon acceptance.

## Supplement 1: A note on sampling depth and the probability of observing zeros

In the metabarcoding literature, it is often asserted that if more reads are sampled, it is more likely that rare variants will be observed. While this is true, the magnitude of the effect is likely less than one would like and only meaningfully changes the probability for a relatively narrow range of rare things. This supplement provides some simple examples for how to do that calculation.

Let us focus on only the last step of the metabarcoding process: multinomial sampling. As we are interested in isolating the contribution of sampling depth we can assert that everything prior to the sampling of DNA strands is identical; only the sampling depth changes. So for illustration, let’s discuss a single taxon (“A”) that comprises 0.0001% of the DNA (1 in 10,000 sequences) of the post-PCR product. We will assume that multinomial sampling is a decent approximation to the process (i.e., there are so many DNA copies floating around removing a few doesn’t materially change the probability of observing a given taxon; for those who are uncomfortable with this assumption, the below can be reframed using the hypergeometric distribution in place of the multinomial). For a single taxon, the multinomial collapses to the binomial (i.e., we can think of many taxa collapsing to two groups: taxon A and not taxon A). The probability mass function for the binomial is:

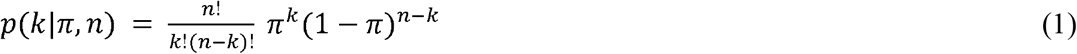

where k is the number of “successes” (observations of taxon A by the sequencer) and *n* is the number of sequences read. We are interested in a single value here: what is the probability of *k=0* (i.e. taxon A was not observed) as the number of sequences examined (*n*) increases. First, simplify for the case *k=0*

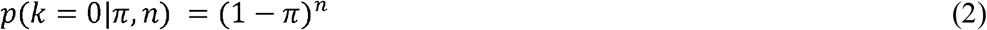

then plug in *π* = 0.0001 a range of values for sampling depth(below I use 10 thousand, 100 thousand, and 1 million reads):

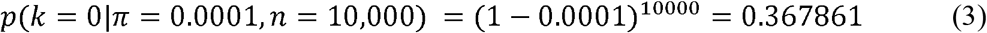

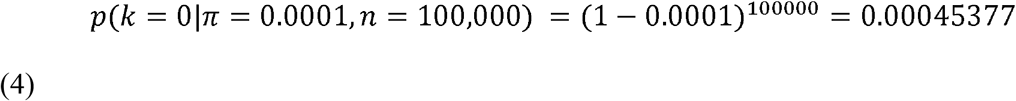

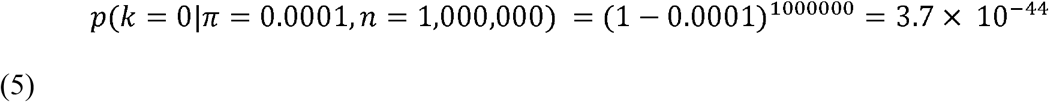

So for a species that is rare we go from seeing 1 or greater sequences with probability of 0.64 (1-0.36) at 10,000 reads to seeing it with almost certainty at 100,000 or more reads. Let’s do the calculation for a rarer sequence.

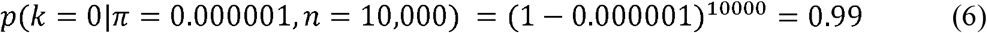

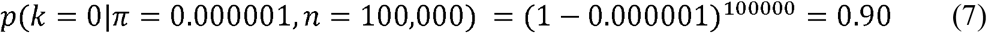

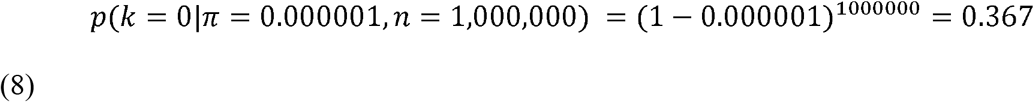

So for a sequence that occurs at a rate of 1 in a million you go from observing 0 99% of the time with a sampling depth of 10,000 reads to 90% at 100 thousand reads to only 36% at 1 million reads.

Going the other direction, let’s look at something that is more common, say 1 in 1,000:

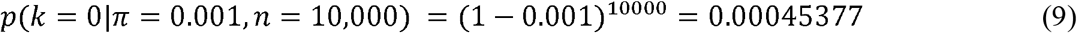

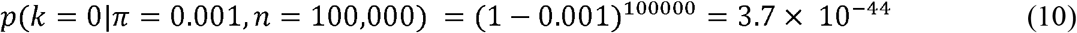

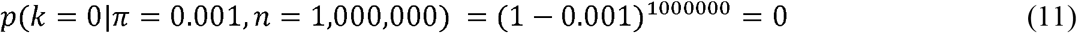

Thus, you are almost certain to see at least one copy at 10,000 or greater read depths.

So what does this mean in general? Basically, you will see rarer things at higher read depths but moving from a read depth of say 10,000 to 1 million will only meaningfully change non-detection of very rare sequence variants (sequences that make up somewhere between 1 in 10,000 and 1 in 1 million copies). If you think that there are a lot of taxa that you care about are in this very rare zone, it may make sense to do more sequencing. But things that are extremely rare (occur at a frequency of less than 1 in a million) still will not be detected. Note that things can be rare after PCR because they are rare in the sample or because they are poor amplifiers, or both. One caveat to the description above is it only includes the probability of observing exactly zero. Many researchers use a higher threshold to determine presence (say *k > 10*, for example). Calculating *k > K* is not quite as easy as *p(k=0)* in that more terms are involved, but it is certainly not a hard calculation and involves summing the probability of *k=0* to *k=K*,

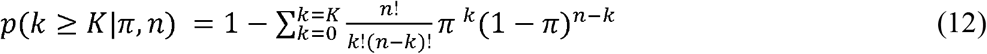

## Supplemental 2 Alternate simulation results

### Changing N_pcr1

It is important to understand how changing some of the parameters in the simulation affect the probability of non-detection. In Fig. S3.1 we used *N_pcr1_* = 20 rather than *N_pcr1_* = 35 presented in the main text. As *N_pcr1_* declines, the probability of non-detects becomes more similar among species and only species with amplification efficiencies that are much lower than the average *a* (in this case *a_i_* < 0.4) have increased non-detection probabilities (Fig. S3.1A).

**Figure S3.1.**
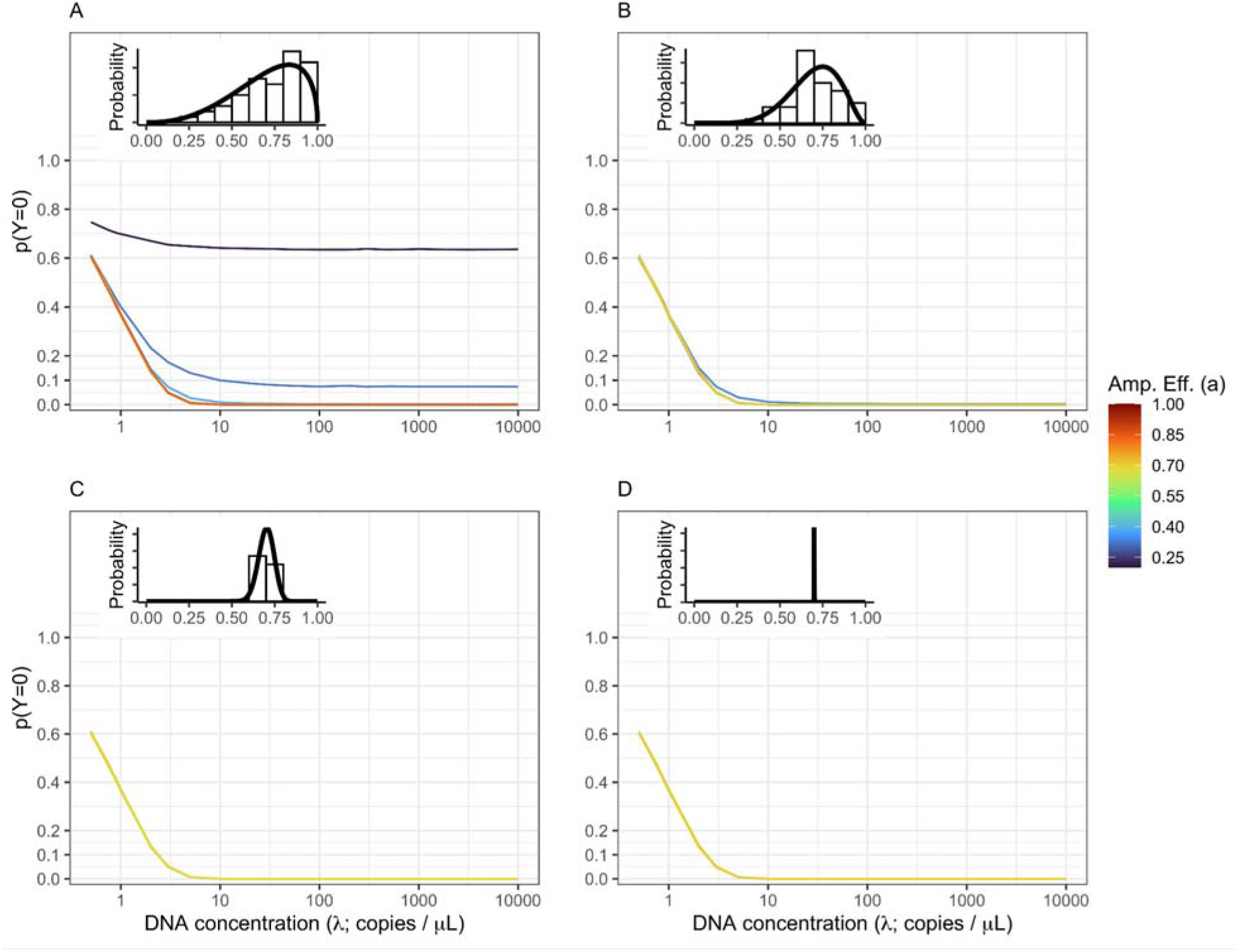
Non-detects Driven By Both DNA Concentration and Amplification Efficiency. The probability of non-detection (*p(Y=0)*) is shown for a community of 50, equally abundant taxa with the amplification efficiency distribution shown inset in each panel. This simulation uses *N_pcr1_* = 20 (see Fig. 1 for the same simulation but with *N_pcr1_* = 35). The amount of among-taxa variation in amplification efficiency varies from highly variable *(A;* γ=5) to moderate variation (*B*: γ=10) to low variation (*C*: γ=100) to effectively no variation (*D*: γ=1,000,000). Both subsampling and amplification efficiencies influence the rate of non-detection. The probability of observing no DNA in a given technical replicate is highest at low DNA concentrations (<10 copies /μL). However, non-detects are possible for species with low amplification efficiencies and very likely (*p(Y=0)* > 0.5) for amplification well below average (in this case approximately *a_i_* < 0.3).

### Simulating uneven DNA concentrations

The base simulation presented in the main text assumes that the starting DNA concentration for each taxon is equivalent (i.e., for ten taxa, each comprises 10% of the DNA in a sample). While this assumption makes it easier to visualize the simulation results, it clearly does not represent natural communities which have skewed abundance distributions (some taxa are common while others are rare). To illustrate the consequences of a skewed abundance distribution we simulated a community of 20 taxa with 2 taxa each comprising 20% of the DNA, 8 taxa each with 5% of the DNA, and 10 taxa with 2% of the DNA. Otherwise, we followed the simulation parameters described in the main text. Figure S3.2 presents the patterns of non-detections for a single community of 20 taxa (Figure S3.2A) with large among-species variation in amplification efficiency (*γ* = 5) and for 20 communities of 20 taxa each overlaid on one figure (Fig. S3.2B). Facets show the true starting proportion within each community (proportions of 0.02, 0.05, or 0.20).

As shown in the even community simulated in the main text, for all taxa non-detection increases as DNA concentration declines and taxa with lower amplification rates show higher probability of non-detection. But there is clearly an interaction between the community proportion and amplification efficiency which affects the probability of non-detection. Specifically, for two taxa with equivalent amplification efficiencies, the more abundant taxa (community proportion of 0.20) have a much lower probability of non-detection than a relatively rare species (community proportion of 0.02; Fig. S3.2B). Indeed, for taxa with a community proportion of 0.02, at a constant DNA concentration, *p*(*Y* = 0|*λ*=10) > 0.5 when *a_i_* < 0.45. In contrast, for taxa with community proportions of 0.20, *p*(*Y* = 0|*λ*=10) > 0.5 only occurred for one taxa in the 20 simulated communities with a very low amplificiation efficiency (*a_i_* = 0.19).

Thus both community proportion and amplification efficiency affect the probability of non-detection. In broad strokes, amplification efficiency will play a more important role in determining non-detection when taxa are rare relative to other species in a sample. The importance of amplification efficiency increases with PCR protocols that use a large number of PCR cycles. Non-detection of relatively common taxa in a community will generally be less influenced by relative amplification efficiency, but non-detection can still occur if amplification efficiency is sufficiently low.

**Figure S3.2.**
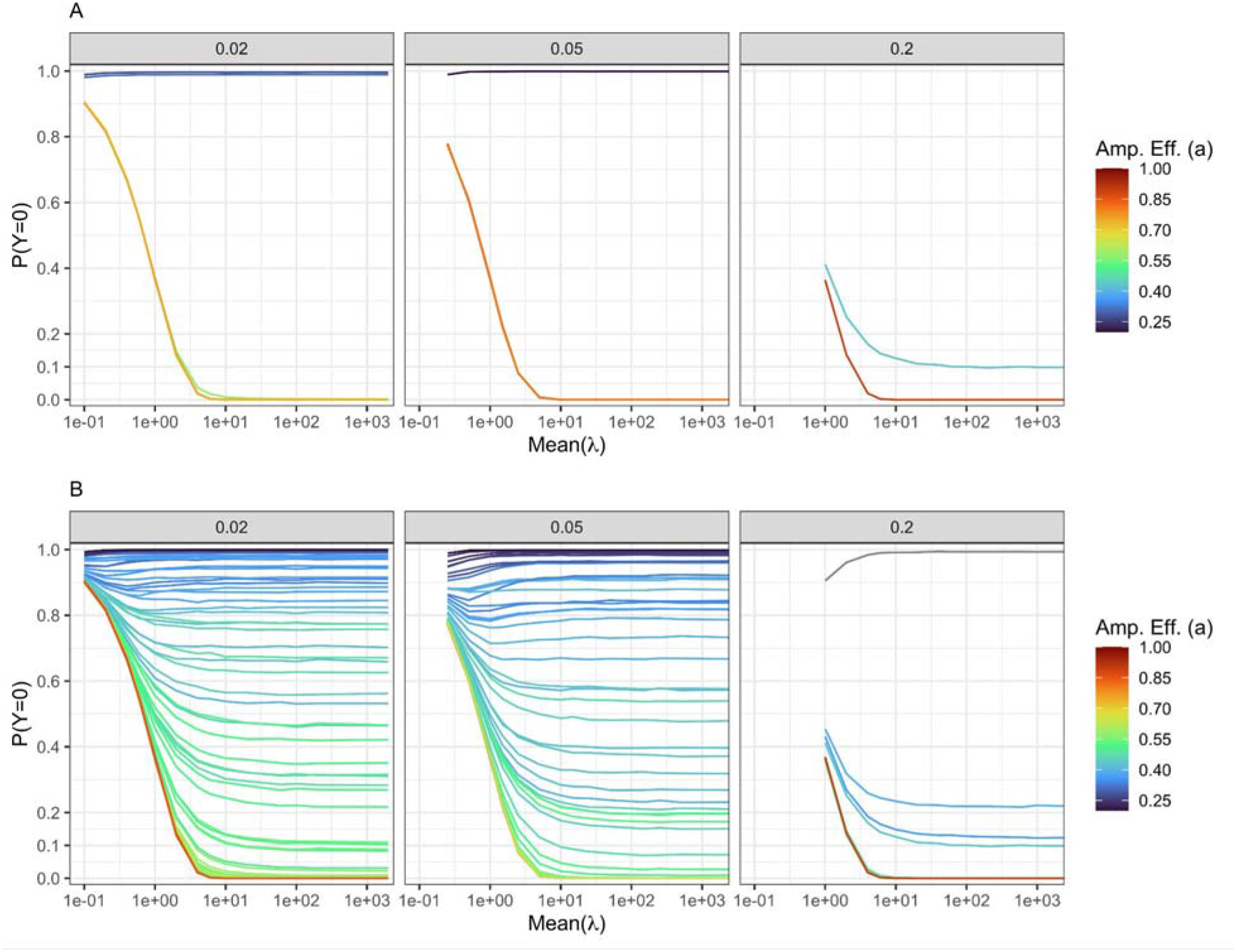
Non-detects Driven By Both DNA Concentration and Amplification Efficiency. The probability of non-detection (*p(Y=0)*) is shown for a community of 20 taxa with 4 taxa comprising 0.20 of the initial DNA, 8 with 0.05 of the DNA, and 10 species comprising 2% of the DNA across a range of initial DNA concentrations. *A*: Presents results for a single 20 taxa community with facets representing the three abundance categories. *B* shows results for 20 communities of 20 taxa each to illustrate general patterns. For all simulations we use *N_pcr1_* = 35 and a fixed amount of among-taxa variation in amplification efficiency (γ = 5). Clearly, relative abundance influence the rate of non-detection with relatively rare taxa (those with 0.02 having larger probabilities of non-detection than common taxa (0.2) with equivalent amplification efficiencies (colors).

